# Scoring Protein Sequence Alignments Using Deep Learning

**DOI:** 10.1101/2021.08.14.456366

**Authors:** Bikash Shrestha, Badri Adhikari

## Abstract

**Background:** A high-quality sequence alignment (SA) is the most important input feature for accurate protein structure prediction. For a protein sequence, there are many methods to generate a SA. However, when given a choice of more than one SA for a protein sequence, there are no methods to predict which SA may lead to more accurate models without actually building the models. In this work, we describe a method to predict the quality of a protein’s SA.

**Methods:** We created our own dataset by generating a variety of SAs for a set of 1,351 representative proteins and investigated various deep learning architectures to predict the local distance difference test (lDDT) scores of distance maps predicted with SAs as the input. These lDDT scores serve as indicators of the quality of the SAs.

**Results:** Using two independent test datasets consisting of CASP13 and CASP14 targets, we show that our method is effective for scoring and ranking SAs when a pool of SAs is available for a protein sequence. With an example, we further discuss that SA selection using our method can lead to improved structure prediction.

## 1 Introduction

All the top-performing groups in the most recent critical assessment of techniques for protein structure prediction (CASP 2020) competition, including Alphafold2 [1], the Baker group [2], the Zhang group [3], and the tFold group [4], have attributed their structure prediction accuracy to the quality of multiple sequence alignments (MSAs or simply SAs in short), generated from the corresponding sequence. This highlights the criticality of SAs for accurate structure prediction. Some groups are investigating if we can predict structures without using evolutionary information. But the accuracy of these methods is significantly lower. Historically, a protein sequence’s SA is understood to be a crucial component in predicting local secondary structures, residue-residue contacts, and distances, and in finding homologous structural templates. Recent methods, however, use raw SA data as an input to directly predict the three-dimensional structures. For example, methods such as Alphafold [5], Alphafold2 [1], trRosetta [6], PrayogRealDist [7], RaptorX [8], tFold [4], and Multicom [9] predict structures or distances using a SA as input.

Although all current protein structure prediction methods use SA generating methods such as HHBlits [10], JackHmmer [11], and DeepMSA [12], there are several ways to generate a variety of SAs for a single protein sequence. For example, AlphaFold uses three databases: UniRef90 [13], BFD [14], and MGnify [15] to obtain related sequences by using HHblits, Jackhammr, and HHsearch to find potential templates. David Baker’s group generated SAs using HHBlits using several rounds of iterative search against the Uniclust [16] database with evalued cutoffs gradually [17]. Additionally, they manually inspected the generated SAs to fine tune the e-value and coverage cutoff in order to increase the metagenomic sequences. Similarly, Tencent lab uses a multi-MSA ensemble [17] approach in their tFold method where they extract 83 groups of SA data from 6 different sequence databases using HHblits, Jackhammr, and PSI-BLAST [18] with different combinations of e-values and iterations. We may observe that all successful methods have their own procedures to generate SAs and it may be difficult to develop a universal method of generating alignments. Also, it has been observed that the choice of a sequence database can significantly affect the quality of the SA generated. For example, in the CASP14 competition, the tFold method from Tencent Lab [4] had an accuracy of 35.13% when they used only one metagenomics database (metaclust50), but after the CASP14 competition, they added the BFD metagenomic database to their method and the accuracy significantly increased to 44.77%. Almost 10% improvement was achieved by adding just one additional database.

The quality of the SA generated determines the accuracy of the predicted structures, particularly in the case of free-modeling or *ab initio* protein sequences. The case of the CASP14 target T1064 serves as an example. For this target, Alphafold2 used a SA generated by their method to build the 3D models but the models generated had low confidence. As a resolution, the group manually selected and added five additional sequences to the SA and re-ran their pipeline. This helped them to achieve more confident models. Similarly, in a recently developed trRosetta method [6], five different alternative alignments were generated using various databases and search algorithms. Four different SAs were generated by searching the Uniclust30 database with HHblits using different e-value cutoffs and by iteratively searching using HHblits followed by Hmmsearch against different metagenomic sequence databases. The optimal SA was selected based on the average probability of the top-L predicted medium and long-range contacts. This SA selection improved the precision of the top-L prediction by 3.1% on the CASP13 FM set and 1.7% on the CAMEO hard targets over their baseline method. Also, in CASP14, the Zhang group generated multiple sets of SAs using variants of their DeepMSA method [12]. The optimal SA was then selected based on the highest cumulative TripletRes [19] probability for the top 10L contacts [17]. These examples further highlight that selecting SA is a key for accurate structure prediction.

Despite this need for a method to rank predicted sequence alignments based on their quality (quality here refers to the usefulness towards building 3D models), no methods have been developed for this purpose. If there existed a method that could select the best alignment for each possible residue pair, a significant improvement of the alignment quality could be gained [20]. Previous studies on SA quality [21, 22, 23] have focused on comparing the performance of SA generation methods using the benchmark databases such as BaliBASE [24], OXBench [25], PREFAB [26], SABmark [27] and IRMBASE [28]. Additionally, [29] describes the use of protein structure predictions to measure the quality of the MSA and [30] describes the use of secondary structure predictions to measure the alignment quality to create benchmarking methods ConTest and QuanTest respectively.

Although protein sequence alignments (SAs) can serve many purposes, in this work, our interest is in predicting the utility of SAs towards accurate structure prediction, i.e., predicting how informative they are for the methods that use them to drive structure prediction. Although it is well understood that the ultimate accuracy of structure prediction depends on many factors other than just the input SA, in general, a SA is the first seed that guides the entire structure prediction process. In this work, we have developed a deep learning-based method to rank multiple sequence alignments based on their quality, i.e., their usefulness for building 3D models. Given a protein sequence and a set of alignments generated for it using multiple methods, our method predicts local distance difference test (lDDT) scores [31] of the distance maps that can be predicted from the SAs. An lDDT score is in the range of [0, 1], where a score of zero indicates a complete mismatch and a score of one suggests identical distance maps. These lDDT scores serve as the alignment quality scores. Irrespective of the method that is used to generate a SA, we predict a score for it. Subsequently, these SAs can be ranked by the scores for selection. Finally, to demonstrate an additional application of our method, we illustrate how it can be used to improve the quality of a SA for 3D structure prediction.

## 2 Methods

### 2.1 Defining SA quality using inter-residue distance lDDT score

We define the quality of a SA as its ‘inform-ability’ for accurate structure prediction. In the absence of a single metric that can be used to assess the quality (utility towards building 3D models) of a SA, we consider the local distance difference test (lDDT) score of the C*β*-distance map, predicted from the SA, as a score that reflects the quality of the MSA. lDDT is a superposition-free score that evaluates differences in distance of all atoms in a model [31]. We use the DISTEVAL tool [32] to calculate the C*β*-lDDT score by ignoring all the pairs below a sequence separation of six residues. For an input SA, we further hypothesize that if we can predict the lDDT score of the inter-residue (inter-C*β*) distance map predicted using the SA, then this score can be used as the SA’s quality. Our hypothesis is based on the findings that accurate inter-residue distance prediction leads to accurate structure prediction [5] [8] [6]. Ideally, one would develop an end-to-end pipeline that builds 3D models (as in AlphaFold2 [1]) and evaluate the accuracy of the models as the quality of the SA. However, developing a fully end-to-end method that predicts 3D models from a SA input is not an easy task (at least as of now). Overall, our formulation of predicting lDDT score as SA quality score serves as to be a useful method for ranking and selecting SAs when we have a multiple choice of MSAs.

### 2.2 Datasets

For training and hyper-parameter optimization of our deep learning models, we curated a representative set of 1351 protein chains obtained from the CATH [33] structural classification database version 4.2. CATH classifies protein structural domains into a hierarchy consisting of 5 levels - class (C), architecture (A), topology/fold (T), superfamily (H), and family. At the topology level, proteins are grouped into fold families depending on the overall shape and connectivity of the secondary structures. From each topology, we selected a protein with the highest length, resulting in 1351 chains. Since we choose a longer protein when possible, around 40% of the proteins in our training sets have *L* = 512 and the remaining are evenly distributed with various lengths. A random subset of 51 chains were used for validation, leaving the remaining for training. We chose the topology level because selecting proteins from a lower level of the hierarchy (for example, superfamily) results in a much larger set requiring computational resources beyond our access.

In addition to the validation set for selecting the deep learning hyper-parameters, we curated two datasets for testing. The first set is SAs predicted by the RaptorX method by the Xu group at the Toyota Technological Institute for 16 CASP13 free-modeling targets. The dataset is available at https://home.ttic.edu/jinbo/. For each target, four SAs are generated with the following parameters: 1) uniclust30 database with e-value 0.001 (uce3), 2) uniclust30 database with e-value 0.00001 (uce5), 3) uniref90 database with e-value 0.001 (ure3), and 4) uniref90 database with e-value 0.00001 (ure5). The second test dataset consists of SAs predicted by the Zhang group at the University of Michigan for 33 protein targets in the CASP14 competition. For each protein target in this dataset, up to 18 SAs are generated using variants of three techniques [3]: using the JGI database containing more than 60 billion microbial genes (mMSA) [34], using the standard DeepMSA pipeline (dMSA), and using a four-stage iterative searching of SAs (qMSA). The original dataset consisted of SAs for 84 targets, but we focused on evaluating only the 33 proteins for which the native (true) structures are available.

### 2.3 Generating SAs for development

To prepare a diverse and representative SA dataset consisting of a wide range of quality, we generated five SAs for each of the 1,351 proteins in our dataset. These SAs were generated using the following techniques: 1) running HHblits [10] at an e-value threshold of 0.0001 (1e-4), coverage of 70%, and iteration set to three (we call these SAs ‘Set A’), 2) e-value threshold of 0.001 (1e-3), coverage of 40%, and iteration set to three (Set B), 3) e-value threshold of 0.1 (1e-1), coverage of 50%, and iteration set to three (Set C), 4) e-value threshold of 10 (1e1), coverage of 30%, and iteration set to one (Set D), and 5) running Jackhmmer [11] with an e-value cutoff at 0.1 (1e-1) with iteration value (n) set to 3 (Set E). We used the uniprot20 (2016_02 version) database with HHblits and uniref90 database with Jackhmmer. Consequently, our training and validation datasets consist of 6500 (1300 x 5) and 255 (51 x 5) examples, respectively. The mean (and standard deviation) of the lDDT scores of the trRosetta generated alignments for the sets D, B, C, A, and E are 0.208 (0.15), 0.218 (0.16), 0.239 (0.13), 0.26 (0.13), and 0.29 (0.18), respectively. Overall, by changing alignment generation parameters such as e-value, coverage, and iteration count, we generated different types of SAs ranging from low to high quality so that our dataset is representative.

### 2.4 Feature generation

For each SA in our training and validation set, we generated a covariance matrix (441 2D channels), amino acid composition (20 1D feature channels), position-specific frequency matrix (21 1D features), and positional entropy (1D feature) where the one dimensional features are translated into 2D (2 x 42 = 84 channels) features by tiling and ‘transposing followed by tiling’ to build input features that can be supplied to a deep learning model. For generating these features, we adapted the scripts in the trRosetta package available at https://github.com/gjoni/trRosetta. We also used the C*β*-distogram predictions made by using trRosetta and a single-channel distance map prediction obtained by ‘flattening’ the distogram as additional input features. A single distogram is a 37-channel tensor where each channel represents the probability of 37 bins. There are 36 bins which are equally spaced with 0.5 Å and one extra bin for distances larger than 20 Å. Flattening here refers to the process of translating a multi-channel distogram into a single channel 2D distance map by computing the average of the lower-bound and upper-bound distances of the range with the highest probability for each residue pair. For example, if the bin [6.5, 7] Å has the highest confidence for a residue pair *i* and *j*, then the flattened distance between *i* and *j* is 6.75 Å. To summarize all of our input features for a protein sequence, we have one SA that translates into a 564 channel volume: 526 channels obtained from the trRosetta scripts, 37 channels from the predicted distogram, and one additional channel from the flattened distogram.

### 2.5 Label generation and deep learning training

For each SA in our dataset, we have a volume (3D features derived from the MSA) as the input and an lDDT score (scalar value) as the output label for training a deep learning model. Specifically, for each SA we first obtain the 564 channel tensor as the input feature and a corresponding lDDT score (calculated by comparing the flattened distance map predicted using trRosetta and a true distance map) as the label for loss calculation during deep learning training. This setting is similar to the image classification problem in computer vision, the difference being a continuous value (regression problem) as an output instead of a class (classification problem). As evaluation metrics, we used Mean absolute error (MAE) and Pearson correlation coefficient (PCC) to calculate the correlations between the lDDT score of the distance map predicted from SA (true label) with the lDDT score predicted by our deep learning model (predicted model output). We started our model architecture search experiments with the traditional VGG16 [35] architecture. However, because of a large model size (number of parameters), the training time per epoch was high, so we switched to testing other architectures. We also experimented with variants of ResNets [36] and also designed our own networks similar to the VGG16 with a smaller number of parameters. None of these networks achieved a PCC higher than 0.8.

Inspired by the recent success of EfficientNets [37], we tested variants of EfficientNets for predicting lDDT scores and observed remarkable improvements in the prediction accuracy. Developers of the EfficientNet architectures have proposed a new compound scaling method that uniformly scales the network width, depth, and resolution. EfficientNets have performed remarkably well in the field of computer vision, achieving much higher accuracy and efficiency than previous ConvNets architectures. In particular, on the ImageNet dataset, EfficientNet-B7 architecture has 84.1% top-1 accuracy, which is a state-of-art performance [37]. Impressively, it is 8.4 times smaller and 6.1 times faster than the previous state-of-the-art ConvNet architectures. In terms of top-1 accuracy, even a much smaller version of EfficientNet, the EfficientNet-B1 architecture, is demonstrated to achieve 79.1% accuracy, outperforming ResNet-152 [36] (77.8%), DenseNet-264 [38] (77.9%), Inception-v3 [39] (78.8%), and Xception [40] (79%). For our model development, we used the smallest version of EfficientNet, known as Efficientnet-B0. We adapted the existing implementation of this network in Keras so that it can accept inputs of different sizes since proteins can be of arbitrary length. We first designed a large model that can accept a protein up to 512 residues long sequences and padded all shorter proteins’ input features with zeros. We call this model M-512. Upon suspecting that such a large model may not perform well for short proteins, we designed three separate models of small, medium, and large size. Specifically, we designed: 1) a model that accepts a SA of a protein sequence up to 128 residues with input ‘shape’ fixed at 128 x 128 x N, i.e, the model can only accept sequences up to 128 residues (M-128), 2) another that accepts up to 256 residues with 256 x 256 x N input (M-256), and 3) another one that accepts up to 512 residues with 512 x 512 x N input (M-512). N is the number of input feature channels, which is equal to 564 here. We also added extra layers at the end of the network to predict a single continuous value. Specifically, we added a ‘Batch Normalization’ layer followed by a 2D convolution layer with a single filter and rectified linear unit (ReLU) as an activation function. As the last layer, we added a ‘Global Average Pooling’ layer to ensure that a single value is predicted regardless of the input size. **Figure 1** summarizes our workflow and the block diagram of our architecture. All networks have 4,231,158 parameters each, regardless of the input size.

**Figure 1:**
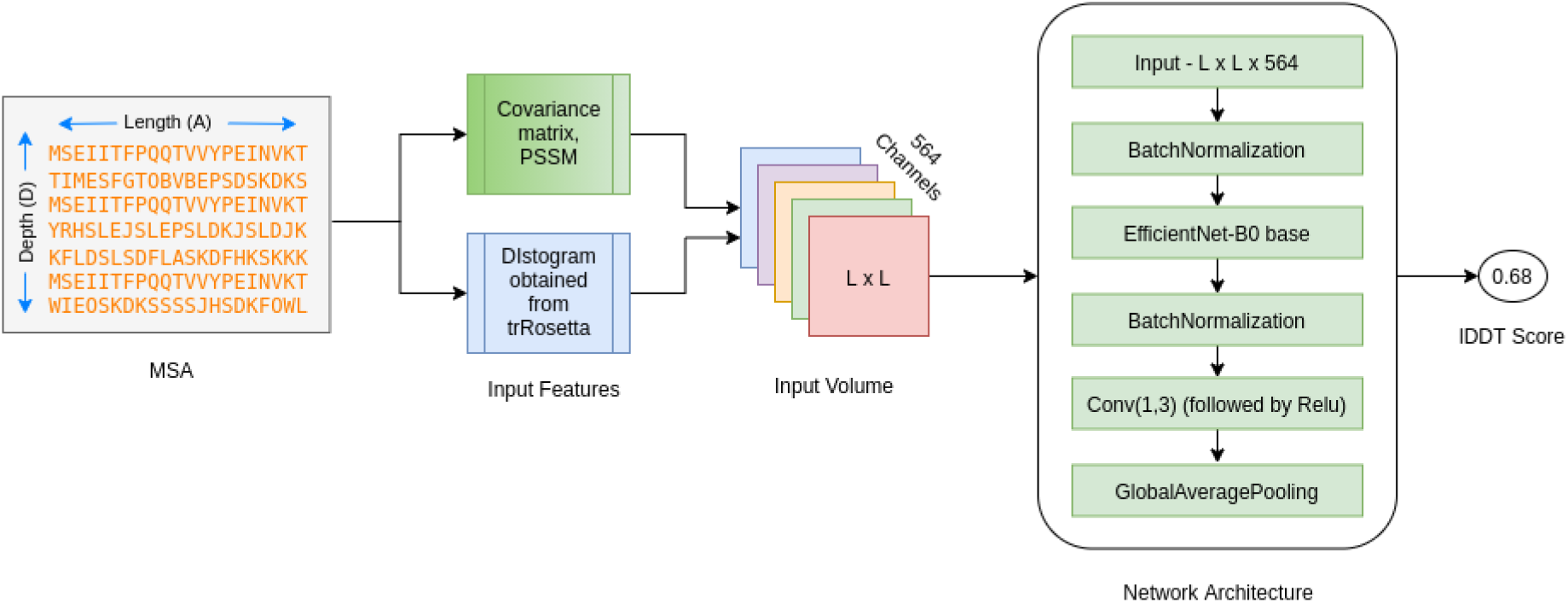
Workflow of our method. The features generated from the SA, including covariance matrix and PSSM, are concatenated with the distogram predictions by trRosetta to create a volume with 564 channels which becomes the input to the deep learning module. The network architecture based on EfficientNet-B0 then predicts an lDDT score.

We describe our training process next. Since our deep learning models are designed to predict an lDDT score, a real number, our supervised learning is a regression problem. We chose mean absolute error (MAE) as the loss function for calculating loss during training. In addition to evaluating the model using MAE, we also calculated the Pearson’s correlation coefficient (PPC) between the true lDDT and predicted lDDT of the validation set as our metric to determine the accuracy of the models. At first, we trained our M-128 model. Since most proteins in our dataset are longer than 128 residues, to train this 128-size model, we chose a 128-length sub-sequence at a random position and calculated the true lDDT score for this sub-sequence as the true label. In other words, for proteins longer than 128 residues, we chop out an input volume of size 128 x 128 x 564 at a random location along the diagonal from the input volume of size L x L x 564, when L > 128. Since we implement this using Keras Data Generators, it also serves as a method for data augmentation. If the size of the protein is smaller than 128, we pad the input volume with zeros to make it 128 x 128 x 564. We trained this model for a total of 128 epochs, monitoring the overall performance using MAE at each epoch and PCC at 16, 32, 48, 64, 96, and 128 epochs. Using a similar approach, we trained the M-256 model. Since a bigger model takes much longer to train, for our M-512 model, we used pre-trained weights from the M-256 and trained only for 8 epochs in order to train faster. For each model, the network weights that deliver the lowest MAE on the validation set are saved as the optimal model. Nesterov-adam (nadam) [41] was chosen as an optimizer.

In a Ubuntu server with dual 4215R 3.2GHz CPUs, 128 GB of RAM, and two NVIDIA Quadro RTX 6000 GPUs, each with 24 GB GPU-memory, one epoch of training the M-256 model takes around 6 hours to complete and M-512 takes around 9.5 hours. All the five sets of input features for our 1353 proteins in the development set occupy around 1.4 terabytes (TB) of space and were stored in solid-state disks (SSDs) for faster access. Python along with the supported libraries Tensorflow, Keras, Scikit-learn, and Numpy were used for development.

## 3 Results

### 3.1 Deep learning model selection using the validation set

In this problem of taking a protein SA as input and predicting an lDDT score, a key challenge is to design a deep learning model that can accept a SA for a protein of arbitrary length and predict a single score. A solution is to design a large model, say for example, a model that can accept proteins up to 512 residues in length and use the model for all protein lengths. To train such a model, input features for all the proteins shorter than 512 residues should be padded. However, such a model could be biased towards predicting the lDDT of longer proteins (close to 512 residues) more accurately. Hence, we designed three models: one that accepts proteins up to 128 residues in length (M-128), one that accepts proteins up to 256 residues (M-256), and finally one up to 512 residues (M-512). We also evaluated the performance when we selected one of the three models at runtime based on the input size (M-Select). Specifically, to generate M-Select prediction for a protein of length *L*, M-128 is selected if *L* ≤ 128, M-256 is selected if 128 < *L* ≤ 256, and M-512 is selected if 256 < *L* ≤ 512. For example, if the input protein length is 120, lDDT prediction from M-128 is chosen as the predicted lDDT. Similarly, if the input length is 450, M-512 is selected for predicting lDDT. Our objective of designing M-128 and M-256 is to check if these models outperform M-512 for shorter proteins.

Our validation dataset consisting of 51 proteins (each with five SAs) was used to investigate which of the four models (M-128, M-256, M-512, or M-Select) performs the best for proteins of all lengths. For this investigation, we predicted lDDT scores for all the 255 (51 x 5) SAs in the validation set using the four models and compared the scores with true lDDT scores. For calculating true lDDT scores, ‘distograms’ predicted by trRosetta were evaluated against the true PDB structures using DISTEVAL [32]. The five SAs for each protein (Set A through Set E) are generated using various alignment generation tools and various parameters for running these tools with an overall goal of generating SAs of diverse quality (see **Methods**).

**Figure 2** shows a comparison of true and predicted lDDT scores for M-128, M-256, M-512, and M-Select, for all five sets of SAs in the 51 validation proteins. It can be observed that the true-vs-prediction scatter plot for M-512 is more accurate than M-128 and M-256. The Pearson’s correlation coefficient (PCC) calculated between true lDDT scores and predicted lDDT scores confirm this observation—PCC values are 0.83, 0.87, and 0.93 for M-128, M-256, and M-512 respectively. Comparison of M-512 and M-Select using PCC values also shows that the two models have similar performance—PCC values are 0.93 and 0.95 for M-512 and M-Select. Since our M-Select method requires all three methods and M-512 is a single model, we chose to use M-512 as our final method for evaluation on other test datasets. The true-vs-prediction plot for M-512 also illustrates that the model is not biased towards longer proteins. Detailed PCC calculations for each of the four models and each of the five SA sets are reported in **Table S1**.

**Figure 2:**
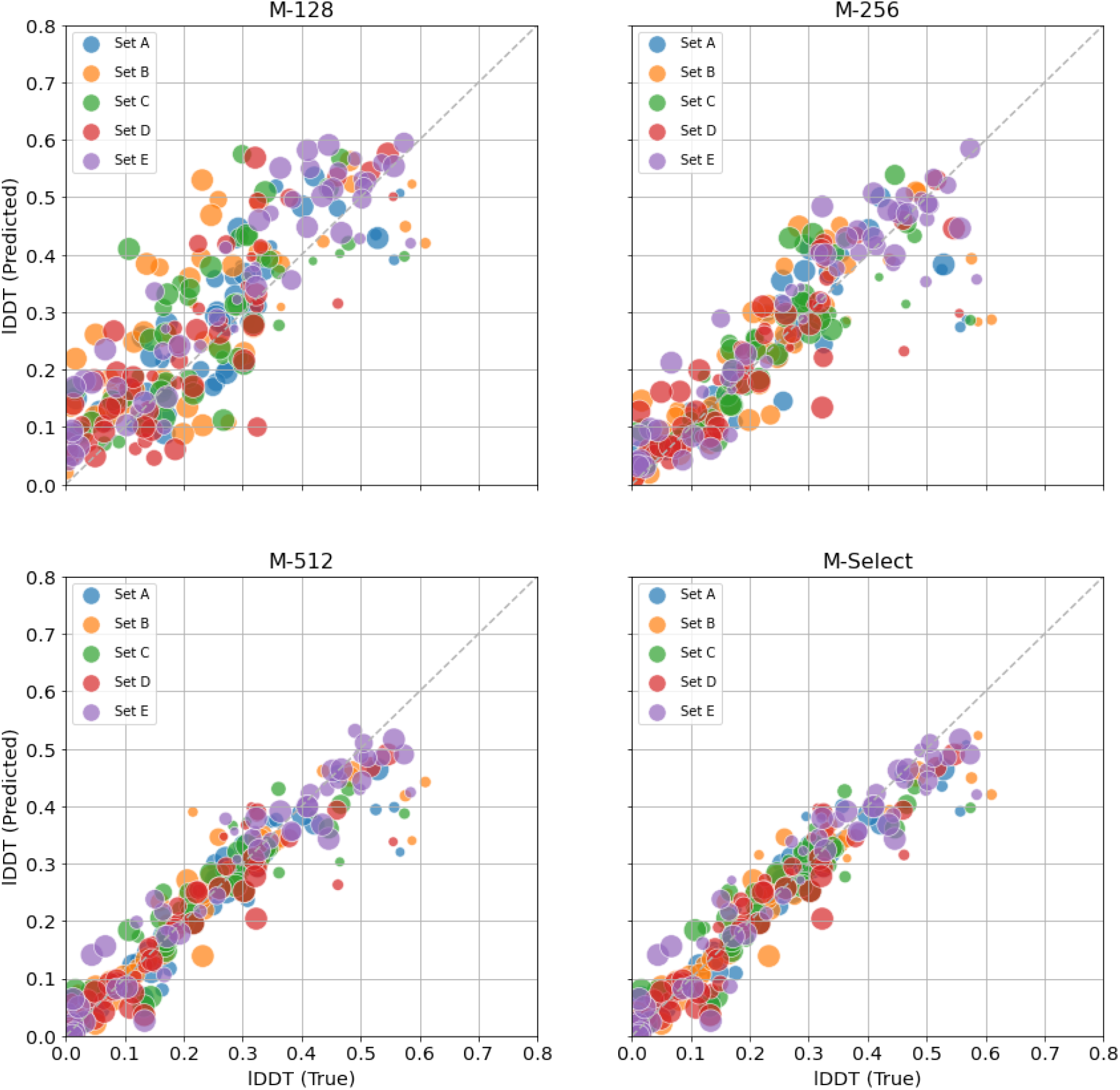
True vs. predicted C*β*-lDDT scores for 51 proteins in the validation set predicted using the four methods: M-128, M-256, M-512, and M-Select. The bubble sizes in all plots correspond to the relative lengths of the proteins, and the color of the points represent the methods used to generate the SAs. Pearson’s correlation coefficient for M-128, M-256, M-512, and M-Select are 0.83, 0.87, 0.93, and 0.95, respectively.

### 3.2 Identifying high-quality SAs via lDDT predictions

Our deep learning model that predicts lDDT score may be used to blindly select a high-quality SA from a pool of SAs. In addition to our validation set, two additional independent datasets are included to discuss our evaluation of SA selection. In our validation set, we have 51 protein sequences, each of which has five different SAs generated (set A through set E) with a variety of quality. Our first independent test set consists of 16 proteins in the free-modeling category of the CASP13 competition, each of which has four SAs. The second independent test set consists of 33 proteins released as targets in the CASP14 competition for which the Zhang group predicted up to 18 SAs. For each SA generated for all of these 100 proteins (51 validation + 16 CASP13 + 33 CASP14), we ran trRosetta to generate input features and ran M-512 to predict lDDT scores. From a pool of SAs for each protein, the SA with the highest predicted lDDT score was selected as the top/best predicted MSA. For evaluation, in each data set, we compare the mean lDDT of our top selected SAs with each individual SA generation method.

**Figure 3**, where we report the summary of our results, shows that for all three datasets, the true mean lDDT scores of the SAs selected by M-512 are higher than any of the individual SA generation methods. For the validation set, the SA generation technique in Set E is best with an average lDDT score of 0.294. Using M-512 to select the best MSA, we obtain a mean lDDT of 0.34. Similarly, for the CASP13 free-modeling set consisting of 16 proteins, SAs selected by M-512 are slightly better (lDDT 0.341 versus 0.337). In this dataset, only a slight improvement is observed over the best SA generation method because of two reasons: 1) the number of choices is limited to four, and 2) the pool does not contain high-quality SAs as the maximum possible lDDT (if we knew the best alignment) is only 0.36. Finally, significant improvements are observed in the case of the CASP14 dataset consisting of 33 proteins. Of the 18 alignment generation methods, the best alignment generation method, ‘qMSA’, has the highest average lDDT of 0.289. When the highest-scored SA is selected based on the M-512 predicted lDDT score, the average lDDT score increases to 0.316. This average lDDT is slightly less than the maximum possible lDDT score of 0.323, i.e, the average lDDT if we knew the best SA for each protein. For all three evaluation datasets, the Pearson’s correlation coefficients (PCC) between the true and predicted lDDT scores for each set of MSAs are higher than 0.8 in most cases. These results are reported in **Table S2**.

**Figure 3:**
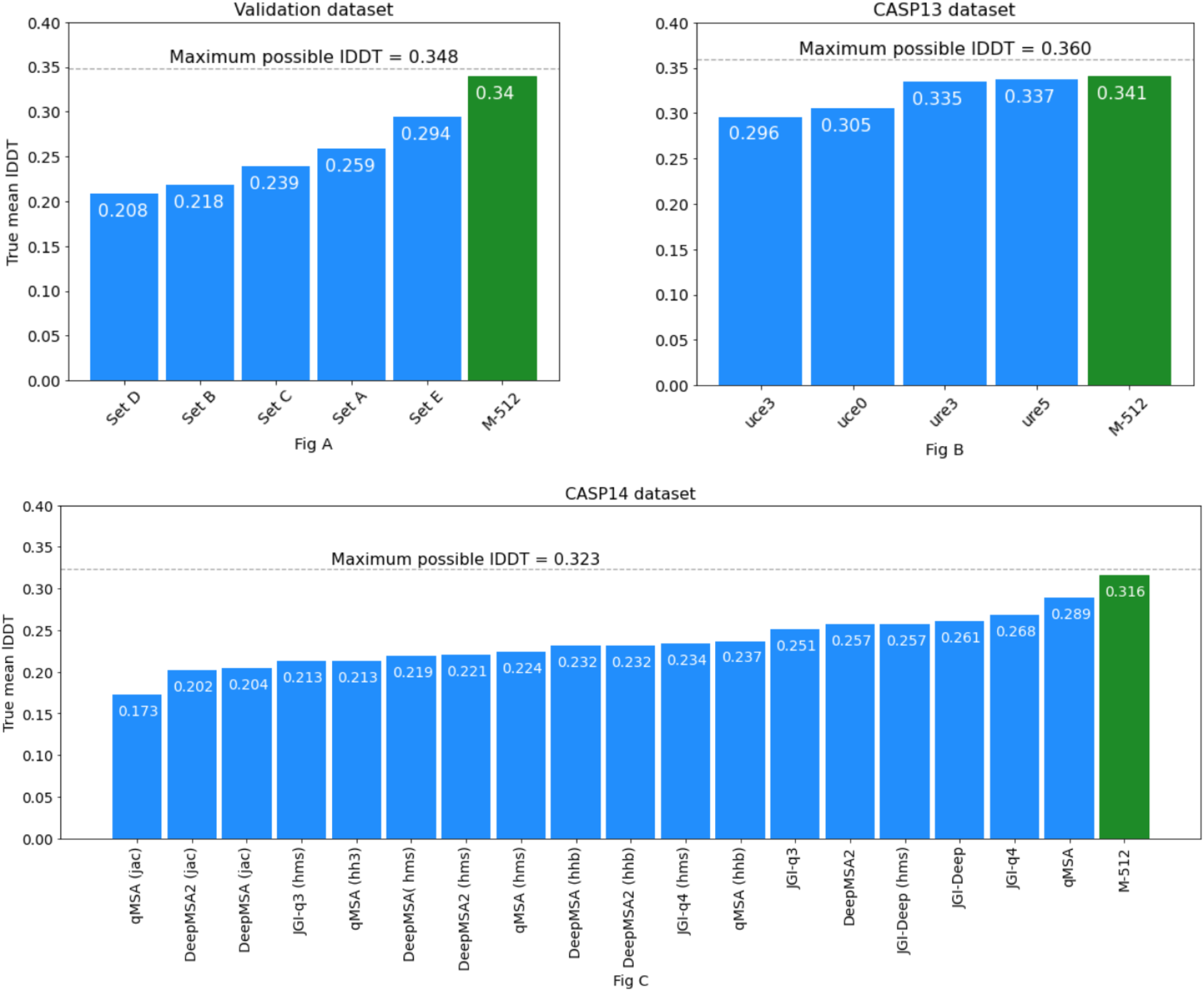
True mean lDDT scores for sequence alignments (SAs) generated using various techniques for the validation dataset (top left plot), the CASP13 FM set (top right plot), and the CASP14 Zhang set (bottom plot). True lDDT scores were obtained by evaluating the trRosetta predicted ‘distograms’ and serve as an indicator of the quality of the overall SA generation method. The mean lDDT score of the top selected SA, using the M-512 model’s scoring, is also reported (green bars). For each dataset, the maximum possible mean lDDT score (the performance of an ideal predictor) is also shown using dotted grey lines.

To investigate why our method is not as effective in selecting the best SA for the CASP13 set as for the validation set and the CASP14 set, we sought to experiment by adding a set of higher quality SAs to the pool of SAs. Specifically, in addition to the four SA sets in the CASP13 set, we added another set of SAs generated in the trRosetta work. These SAs are accessible at https://yanglab.nankai.edu.cn/trRosetta/. The average true lDDT of the distogram predictions obtained from these SAs is 0.437, which is much higher than for any of the four sets (mean lDDT scores with 0.305, 0.296, 0.335, and 0.337). After adding this SA set, we re-ran our M-512 model. The average true lDDT score of the SA selected by M-512 is 0.439, which is slightly higher than the trRosetta set. Of the 16 proteins, M-512 correctly selected the best SA for 14 proteins.

To further cross-check our method of SA ranking and selection, for the 33 protein sequences in the Zhang CASP14 set, we compared our method with Zhang group’s SA selection and ranking presented in the CASP14 conference by the group [3]. The Zhang dataset also includes a SA named ‘protein.a3m’ that the group selected as the final alignment for distance and structure prediction. For each protein, the Zhang group had scored and ranked SAs using their in-house method that selects the best SA among the 18 SAs based on the highest cumulative TripletRes [19] predicted probability for the top 10 L contacts [17]. Here, L refers to the target sequence length. The average lDDT score of these 33 SA files (‘protein.a3m’ files) selected by the Zhang method is 0.312. For calculating this lDDT score, we used the alignment as input to the trRosetta pipeline and evaluated predicted distances using DISTEVAL, i.e., the same process used for calculating all other lDDT scores in this work. This lDDT score of 0.312 can be viewed as a benchmark for assessing our methods. For the same set, ranking and selecting SA using our M-512 method, on the other hand, we obtain an average lDDT score of 0.316. These results suggest that our M-512 method performs on par or better than the Zhang group’s SA scoring and selection method.

### 3.3 An alternative method for feature generation

For our experiments, we chose to use the features and distogram predictions from trRosetta, but there are alternatives. For example, the ProSPr [42] method also runs entirely on Tensorflow and predicts distograms. To fully validate that our method development is independent of the choice of a feature generation tool, we would need to generate the features using an alternative method such as ProSPr and repeat our training and evaluation. However, if the lDDT scores predicted by trRosetta and ProSPr are the same (or close) then repeating the training experiments can be considered redundant. Hence, we sought to compare the lDDT scores predicted by ProSPr and trRosetta using the same input SAs. For this comparison, we chose the CASP14 dataset consisting of 33 proteins and 18 sets of alignments. On this dataset, for the 18 SA sets, the Pearson’s correlation coefficient (PCC) between lDDT scores calculated from distograms predicted by trRosetta and by ProSPr range from 0.92 to 0.97. The overall PCC between all the lDDT scores from trRosetta and ProSPr distances is 0.95. The comparison of lDDT scores in **Figure 4** shows that the lDDT scores for ProSPr are slightly lower than that of trRosetta. One possible reason for the lower lDDT of ProSPr predicted distograms is because the distograms by ProSPr have bigger bins of 2 Å range (0-4 Å, 4-6 Å, 6-8 Å, etc.) compared to more granular trRosetta’s bins of 0.5 Å (2-2.5 Å, 2.5-3 Å, 3-3.5 Å, etc). During the flattening process, smaller bins translate to higher resolution distances, causing the flattened distance map to yield a higher lDDT score.

**Figure 4:**
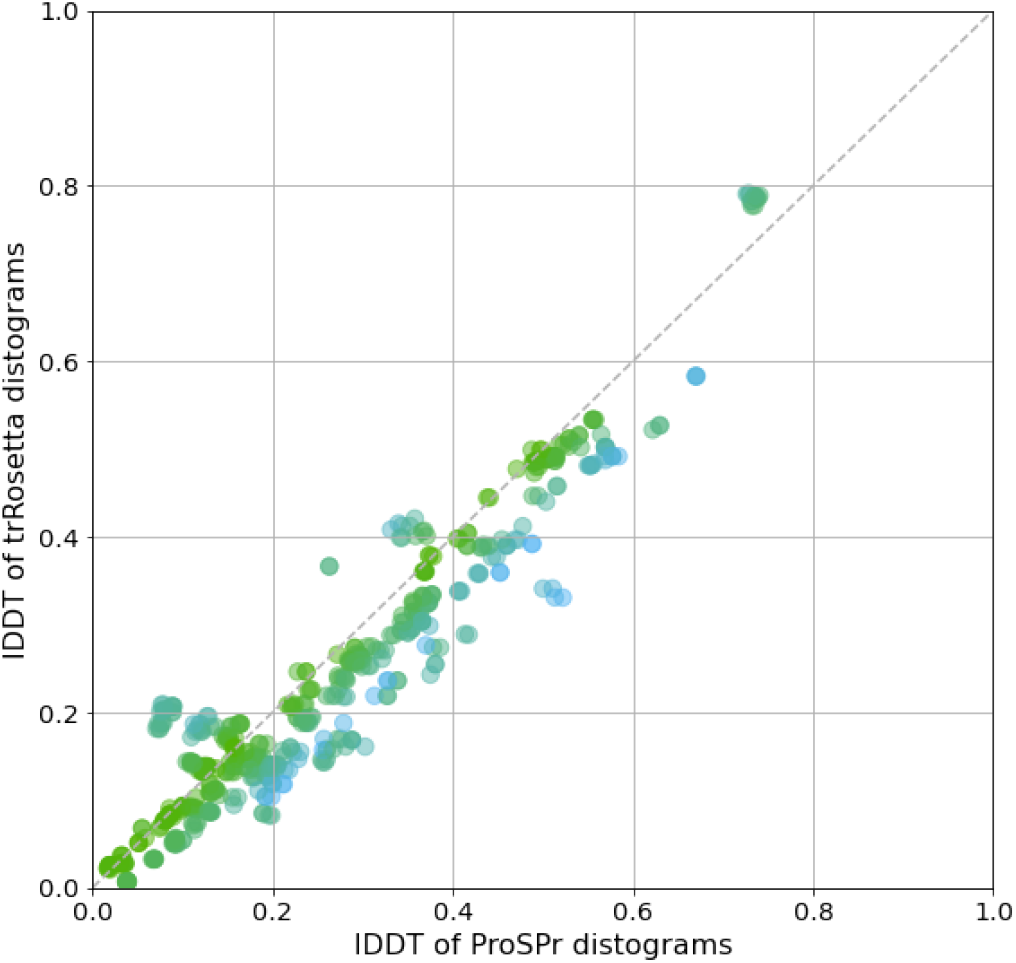
Comparison of lDDT scores of distograms predicted by trRosetta and ProSPr for the 33 targets (times 18 SA combinations) in the CASP14 dataset. Pearson’s correlation coefficient between the two sets is 0.95.

### 3.4 Selecting SA leads to more accurate structure prediction

In this section, we discuss the application of SA selection to build more accurate protein three-dimensional (3D) models. From our CASP14 dataset, we selected all single-domain targets with less than 250 residues (*L* ≤ 250) and built 3D models using DISTEVAL [32], which extends CONFOLD [43] to accept distance restraints instead of contacts. For these 18 targets, we built 3D models with SAs from the the best alignment generation method ‘qMSA’. The average TM-score of the top-one model (out of 20 model decoys), in this case, is 0.388. On the other hand, the average TM-score for the top-one model generated using the SAs selected by M-512 is 0.401, with improvement observed in 13 out of 18 cases. Our model building method, however, is naive and the decoy size is only a handful (20 models). Hence, our results only serve as proof of the concept and improvements should be much more remarkable if more powerful methods such as Rosetta [44] or i-Tasser [45] are used. Detailed results are reported in **Table S3**. Finally, with a CASP14 free-modeling (hard) target T1041 (*L* = 242) as an example, next we discuss how improved SA selection can lead to improved structure prediction. For this target, the Zhang group had a total of twelve SA predictions using their DeepMSA2 method and additional six SAs using the JGI database—18 SA options in total. These SAs yield a variety of true lDDT scores ranging from 0.007 to 0.46. The TM-score of the top-one model (out of 20) with ‘qMSA’ SA as the input is 0.62. The ‘qMSA’ method of alignment generation is the best of all 18 methods. Prediction of lDDT scores using our method M-512 and ranking, enabled us to precisely select the best SA as our top one. The TM-score of the top model for this top-ranked SA is 0.71 (see **Figure S1**). This example demonstrates that SA selection can be effectively used to select high-quality SAs from a pool of SAs and can ultimately help in building more accurate 3D models.

### 3.5 Bootstrapping to improve SA quality

As an additional example application of our method in the field of structure prediction, we investigated if the quality of a SA can be improved even in the absence of a native structure for the input protein sequence. A sequence alignment constitutes of aligned sequences and each of them contribute to the overall quality of the SA. When building a SA via remote sequence-homology search, poor quality sequences also get collected. This follows that if we bootstrap, i.e., randomly sub-sample sequences from a SA, and create a new SA, there is a probability that such a sub-SA has a higher quality. If we generate many such sub-SAs, some of these decoy sub-SAs may be better in quality than the original SA. To test this hypothesis, we randomly selected 15 protein targets within the CASP13 set and generated 100 sub-SAs for each of the four SA sets (uce0, uce3, ure3, and ure5), by removing up to 40% of the sequences selected randomly. These 100 decoy SAs were then passed through our M-512 model to predict the lDDT scores and the SAs within the SA set were ranked using the predicted scores. Finally, for each SA set, we compared the true lDDT score of the top-ranked sub-SA as well as the best sub-SA among the top-5 sub-SAs (ranked by predicted lDDT) with the true lDDT of the original SA. Our results show that we could obtain a higher quality SA for 49 out of 60 cases with up to 12.68% improvement in lDDT score (see **Table S4**). These results suggest that, in future, advanced methods of bootstrapping and generating more decoys will likely yield significantly better results.

## 4 Conclusion

In this work, we demonstrated that protein sequence alignments (SAs) can be scored with the help of deep learning without building three-dimensional models. We also illustrated that selecting a high-quality SA from a pool of SAs can lead to more accurate structure predictions. Our findings suggest that deep learning can be useful for not just selecting an SA from a pool but also to sieve poor quality sequences out from a given SA. We also speculate that using more recent deep learning architectures such as attention networks can also lead to more accurate results.

## 5 Availability

Our methods and data are available at https://github.com/ba-lab/Alignment-Score/.

## 6 Acknowledgement

We are thankful for the financial support received from the US National Science Foundation to B.A. (award number 1948117). We would also like to thank Dr. Jinbo Xu at the Toyota Technological Institute for making the CASP13 sequence alignment data public. We are grateful to Dr. Yang Zhang at the University of Michigan and Dr. Chengxing Zhang, currently at Yale University, for sharing their CASP14 sequence alignment data. We are thankful to Dr. Jie Hou at Saint Louis University, Dr. Sebastian Bittrich at Research Collaboratory for Structural Bioinformatics (RCSB) PDB, and Dr. Dennis Della Corte at Brigham Young University for their invaluable comments and suggestions to improve our manuscript. Last but not the least, we are thankful to graduate students Navneet Kaur, Amulya Reddy Lakku, and James Smith at the University of Missouri-St. Louis and Emma Scally at Mary Institute Country Day School for proofreading our manuscript and suggesting necessary corrections.

## Notes

https://github.com/ba-lab/Alignment-Score/

### Competing Interest Statement

The authors have declared no competing interest.

## References

[1] Jumper, J., Evans, R., Pritzel, A., Green, T., Figurnov, M., Tunyasuvunakool, K., Ronneberger, O., Bates, R., Žídek, A., Bridgland, A., Meyer, C., A A Kohl, S., Potapenko, A., J Ballard, A., Cowie, A., Romera-Paredes, B., Nikolov, S., Jain, R., Adler, J., Back, T., Petersen, S., Reiman, D., Steineggerù, M., Pacholska, M., Silver, D., Vinyals, O., W Senior, A., Kavukcuoglu, K., Kohli, P., and Hassabis, D. (2020) 2020 CASP14 Conference., p. 42.

[2] Anishchenko, I., Baek, M., Park, H., Dauparas, J., Hiranuma, N., Mansoor, S., Humphrey, I., and Baker, D. (2020) Tertiary structure (TS) prediction and refinement from Baker groups., p. 33.

[3] Zheng, W., Li, Y., Zhang, C., and Zhou, X. (2020) Integration of threading and deep learning for protein structure prediction., p. 21.

[4] Wang, S., Lan, H., Shen, T., Wu, J., Zheng, L., Pei, J., Liu, Y., Huang, J., Huang, N., Xu, Z., Liu, W., and Huang, J. (2020) Accurate Contact/Distance Prediction by tFold., 24.

[5] Senior, A. W., Evans, R., Jumper, J., Kirkpatrick, J., Sifre, L., Green, T., Qin, C., Žídek, A., Nelson, A. W., Bridgland, A., et al. (2020) Improved protein structure prediction using potentials from deep learning. Nature, 577(7792), 706–710.

[6] Yang, J., Anishchenko, I., Park, H., Peng, Z., Ovchinnikov, S., and Baker, D. (2020) Improved protein structure prediction using predicted interresidue orientations. Proceedings of the National Academy of Sciences, 117(3), 1496–1503.

[7] Adhikari, B. (2020) REALDIST: Real-valued protein distance prediction. bioRxiv,.

[8] Xu, J. (2019) Distance-based protein folding powered by deep learning. Proceedings of the National Academy of Sciences, 116(34), 16856–16865.

[9] Liu, J., Wu, T., Guo, Z., Hou, J., and Cheng, J. (2021) Improving protein tertiary structure prediction by deep learning and distance prediction in CASP14. bioRxiv,.

[10] Remmert, M., Biegert, A., Hauser, A., and Söding, J. (2012) HHblits: lightning-fast iterative protein sequence searching by HMM-HMM alignment. Nature methods, 9(2), 173–175.

[11] Johnson, L. S., Eddy, S. R., and Portugaly, E. (2010) Hidden Markov model speed heuristic and iterative HMM search procedure. BMC bioinformatics, 11(1), 1–8.

[12] Zhang, C., Zheng, W., Mortuza, S., Li, Y., and Zhang, Y. (2020) DeepMSA: constructing deep multiple sequence alignment to improve contact prediction and fold-recognition for distant-homology proteins. Bioinformatics, 36(7), 2105–2112.

[13] Consortium, U. (2019) UniProt: a worldwide hub of protein knowledge. Nucleic acids research, 47(D1), D506–D515.

[14] Steinegger, M., Mirdita, M., and Söding, J. (2019) Protein-level assembly increases protein sequence recovery from metagenomic samples manyfold. Nature methods, 16(7), 603–606.

[15] Mitchell, A. L., Almeida, A., Beracochea, M., Boland, M., Burgin, J., Cochrane, G., Crusoe, M. R., Kale, V., Potter, S. C., Richardson, L. J., et al. (2020) MGnify: the microbiome analysis resource in 2020. Nucleic acids research, 48(D1), D570–D578.

[16] Mirdita, M., von den Driesch, L., Galiez, C., Martin, M. J., Söding, J., and Steinegger, M. (2017) Uniclust databases of clustered and deeply annotated protein sequences and alignments. Nucleic acids research, 45(D1), D170–D176.

[17] CASP14 (2020) CRITICAL ASSESSMENT OF TECHNIQUES FOR PROTEIN STRUCTURE PREDICTION., 345.

[18] Altschul, S. F., Madden, T. L., Schäffer, A. A., Zhang, J., Zhang, Z., Miller, W., and Lipman, D. J. (1997) Gapped BLAST and PSI-BLAST: a new generation of protein database search programs. Nucleic acids research, 25(17), 3389–3402.

[19] Li, Y., Zhang, C., Bell, E. W., Zheng, W., Zhou, X., Yu, D.-J., and Zhang, Y. (2021) Deducing high-accuracy protein contact-maps from a triplet of coevolutionary matrices through deep residual convolutional networks. PLoS computational biology, 17(3), e1008865.

[20] Elofsson, A. (2002) A study on protein sequence alignment quality. Proteins: Structure, Function, and Bioinformatics, 46(3), 330–339.

[21] Ahola, V., Aittokallio, T., Vihinen, M., and Uusipaikka, E. (2006) A statistical score for assessing the quality of multiple sequence alignments. BMC bioinformatics, 7(1), 1–19.

[22] Aniba, M. R., Poch, O., and Thompson, J. D. (2010) Issues in bioinformatics benchmarking: the case study of multiple sequence alignment. Nucleic acids research, 38(21), 7353–7363.

[23] Wang, Y., Wu, H., and Cai, Y. (2018) A benchmark study of sequence alignment methods for protein clustering. BMC bioinformatics, 19(19), 95–104.

[24] Thompson, J. D., Koehl, P., Ripp, R., and Poch, O. (2005) BAliBASE 3.0: latest developments of the multiple sequence alignment benchmark. Proteins: Structure, Function, and Bioinformatics, 61(1), 127–136.

[25] Raghava, G., Searle, S. M., Audley, P. C., Barber, J. D., and Barton, G. J. (2003) OXBench: a benchmark for evaluation of protein multiple sequence alignment accuracy. BMC bioinformatics, 4(1), 1–23.

[26] Edgar, R. C. (2004) MUSCLE: multiple sequence alignment with high accuracy and high throughput. Nucleic acids research, 32(5), 1792–1797.

[27] Van Walle, I., Lasters, I., and Wyns, L. (2005) SABmark—a benchmark for sequence alignment that covers the entire known fold space. Bioinformatics, 21(7), 1267–1268.

[28] Subramanian, A. R., Weyer-Menkhoff, J., Kaufmann, M., and Morgenstern, B. (2005) DIALIGN-T: an improved algorithm for segment-based multiple sequence alignment. BMC bioinformatics, 6(1), 1–13.

[29] Fox, G., Sievers, F., and Higgins, D. G. (2016) Using de novo protein structure predictions to measure the quality of very large multiple sequence alignments. Bioinformatics, 32(6), 814–820.

[30] Le, Q., Sievers, F., and Higgins, D. G. (2017) Protein multiple sequence alignment benchmarking through secondary structure prediction. Bioinformatics, 33(9), 1331–1337.

[31] Mariani, V., Biasini, M., Barbato, A., and Schwede, T. (2013) lDDT: a local superposition-free score for comparing protein structures and models using distance difference tests. Bioinformatics, 29(21), 2722–2728.

[32] Adhikari, B., Shrestha, B., Bernardini, M., Hou, J., and Lea, J. (2021) DISTEVAL: a web server for evaluating predicted protein distances. BMC bioinformatics, 22(1), 1–9.

[33] Pearl, F. M., Bennett, C., Bray, J. E., Harrison, A. P., Martin, N., Shepherd, A., Sillitoe, I., Thornton, J., and Orengo, C. A. (2003) The CATH database: an extended protein family resource for structural and functional genomics. Nucleic acids research, 31(1), 452–455.

[34] Chen, I.-M. A., Chu, K., Palaniappan, K., Ratner, A., Huang, J., Huntemann, M., Hajek, P., Ritter, S., Varghese, N., Seshadri, R., et al. (2021) The IMG/M data management and analysis system v. 6.0: new tools and advanced capabilities. Nucleic Acids Research, 49(D1), D751–D763.

[35] Simonyan, K. and Zisserman, A. (2014) Very deep convolutional networks for large-scale image recognition. arXiv preprint arXiv:1409.1556,.

[36] He, K., Zhang, X., Ren, S., and Sun, J. (2016) Deep residual learning for image recognition. In Proceedings of the IEEE conference on computer vision and pattern recognition pp. 770–778.

[37] Tan, M. and Le, Q. (2019) Efficientnet: Rethinking model scaling for convolutional neural networks., pp. 6105–6114.

[38] Huang, G., Liu, Z., Van Der Maaten, L., and Weinberger, K. Q. (2017) Densely connected convolutional networks., pp. 4700–4708.

[39] Szegedy, C., Vanhoucke, V., Ioffe, S., Shlens, J., and Wojna, Z. (2016) Rethinking the inception architecture for computer vision., pp. 2818–2826.

[40] Chollet, F. (2017) Xception: Deep learning with depthwise separable convolutions., pp. 1251–1258.

[41] Dozat, T. (2016) Incorporating nesterov momentum into adam.,.

[42] Billings, W. M., Hedelius, B., Millecam, T., Wingate, D., and Della Corte, D. (2019) ProSPr: democratized implementation of alphafold protein distance prediction network. BioRxiv, p. 830273.

[43] Adhikari, B., Bhattacharya, D., Cao, R., and Cheng, J. (2015) CONFOLD: residue-residue contact-guided ab initio protein folding. Proteins: Structure, Function, and Bioinformatics, 83(8), 1436–1449.

[44] Park, H., Lee, G. R., Kim, D. E., Anishchenko, I., Cong, Q., and Baker, D. (2019) High-accuracy refinement using Rosetta in CASP13. Proteins: Structure, Function, and Bioinformatics, 87(12), 1276–1282.

[45] Yang, J., Yan, R., Roy, A., Xu, D., Poisson, J., and Zhang, Y. (2015) The I-TASSER Suite: protein structure and function prediction. Nature methods, 12(1), 7–8.

