## Supplemental Document (PDF) for "Scoring Protein Sequence Alignments Using Deep Learning"

Table **S1**: Each model's prediction was compared to the true score. The given correlations are calculated between the true lddt scores and predicted lddt scores for each combination of the validation set. Pearson correlation coefficient for all the three models along with combined for the validation set are given in the table. Five MSAs combinations are represented as Set 1, Set 2, Set 3 and so on respectively.

| Model | PCC between true and predicted lDDT scores | | | | | |
|---|---|---|---|---|---|---|
| | Set A | Set B | Set C | Set D | Set E | All 5 sets |
| M-128 | 0.803 | 0.725 | 0.811 | 0.808 | 0.887 | 0.827 |
| M-256 | 0.814 | 0.822 | 0.851 | 0.859 | 0.929 | 0.869 |
| M-512 | 0.916 | 0.915 | 0.911 | 0.919 | 0.958 | 0.930 |
| M-Select | 0.951 | 0.947 | 0.931 | 0.948 | 0.957 | 0.950 |

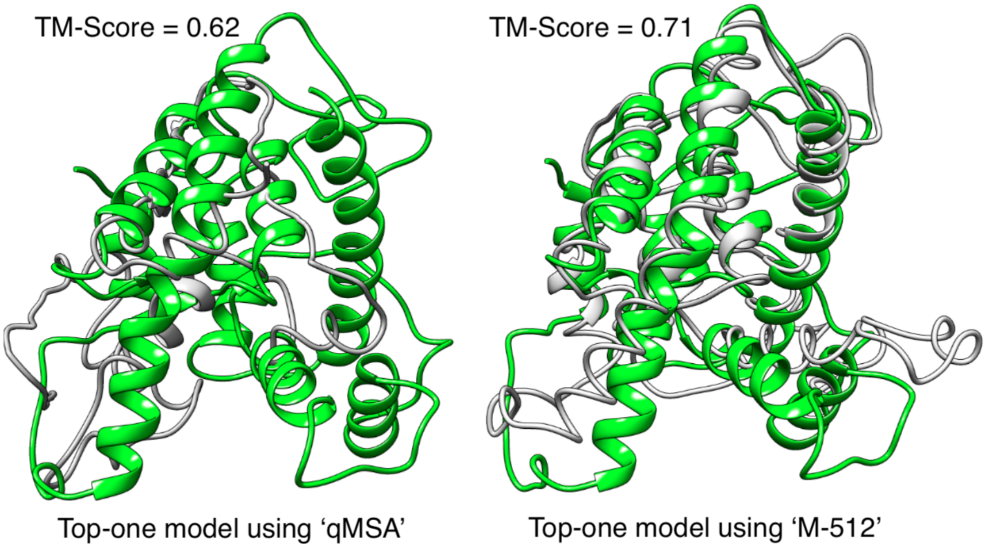


Figure **S1**: Comparison of top-one model (out of 20) shown in grey with the corresponding true structure shown in green for the CASP14 target T1041 with input alignments from the best SA generation method 'qMSA' (on the left) and from the SA selected using our method M-512 (on the right). Models were reconstructed using DISTFOLD and visualized using UCSF Chimera.

Table **S2**: True mean lDDT scores (second last column) and Pearson's correlation coefficient (PCC) between true and predicted lDDT scores (last column) for multiple sequence alignments (MSAs) generated using various techniques for the validation dataset and two other independent test datasets. True lDDT scores were obtained by evaluating the trRosetta predicted 'distograms' and serve as an indicator of the quality of the overall MSA generation method. For each MSA set, the mean lDDT score of the top selected MSA, using the M-512 model's scoring, is also reported. For each dataset, the maximum possible mean lDDT score, i.e., the performance of an ideal predictor, is reported too.

| Dataset | Method for building MSA | True Mean lDDT | PCC |
|---|---|---|---|
| , | Set A | 0.259 | 0.92 |
| | Set B | 0.218 | 0.92 |
| | Set C | 0.239 | 0.91 |
| Cath validation set | Set D | 0.208 | 0.92 |
| | Set E | 0.294 | 0.96 |
| | Maximum possible lDDT | 0.348 | N/A |
| | MSA selected using M-512 | 0.340 | N/A |
| | uce0 | 0.305 | 0.82 |
| | uce3 | 0.296 | 0.93 |
| CASP13 FM RaptorX set | ure3 | 0.335 | 0.91 |
| | ure5 | 0.337 | 0.89 |
| | Maximum possible lDDT | 0.360 | N/A |
| | MSA selected using M-512 | 0.341 | N/A |
| | DeepMSA2 (jac) | 0.202 | 0.82 |
| | DeepMSA (jac) | 0.204 | 0.85 |
| CASP14 Zhang set | DeepMSA (hms) | 0.219 | 0.82 |
| | DeepMSA2 (hms) | 0.221 | 0.86 |
| | DeepMSA2 (hhb) | 0.232 | 0.86 |
| | DeepMSA | 0.232 | 0.86 |
| | DeepMSA2 | 0.257 | 0.89 |
| | JGI-q3 | 0.251 | 0.79 |
| | JGI-Deep (hms) | 0.257 | 0.80 |
| | JGI-Deep | 0.261 | 0.80 |
| | JGI-q4 | 0.268 | 0.81 |
| | JGI-q3 (hms) | 0.213 | 0.839 |
| | JGI-q4 (hms) | 0.234 | 0.887 |
| | qMSA | 0.289 | 0.83 |
| | qMSA (hms) | 0.224 | 0.901 |
| | qMSA (hhb) | 0.237 | 0.839 |
| | qMSA (hh3) | 0.213 | 0.901 |
| | qMSA (jac) | 0.173 | 0.893 |
| | Maximum possible lDDT | 0.360 | N/A |
| | MSA selected using M-512 | 0.341 | N/A |

Table **S3**: Single domain targets shorter than 250 residues in the CASP14 dataset, along with TM-score of the top-one model for models built using the MSA from best alignment generation method 'qMSA' and the MSA ranked on top by our method M-512.

| Model | Length | TM-Score of top-1 model | | Top-ranked SA (using M-512) |
|---|---|---|---|---|
| | | qMSA as input | Top-ranked SA as input | |
| T1026 | 172 | 0.360 | 0.307 | DeepMSA2.hhba3m |
| T1029 | 125 | 0.499 | 0.468 | q3JGI.a3m |
| T1031 | 95 | 0.323 | 0.480 | q4JGI.a3m |
| T1033 | 100 | 0.224 | 0.228 | DeepMSA2 |
| T1035 | 102 | 0.701 | 0.759 | q4JGI.a3m |
| T1038 | 199 | 0.247 | 0.312 | DeepMSA2 |
| T1039 | 161 | 0.346 | 0.332 | q4JGI.a3m |
| T1040 | 130 | 0.260 | 0.342 | q4JGI.a3m |
| T1041 | 242 | 0.624 | 0.710 | q4JGI.a3m |
| T1043 | 148 | 0.153 | 0.154 | q3JGI.a3m |
| T1046s1 | 74 | 0.677 | 0.677 | qMSA |
| T1046s2 | 142 | 0.298 | 0.407 | DeepJGI.hmsa3m |
| T1049 | 141 | 0.557 | 0.259 | q3JGI.a3m |
| T1054 | 190 | 0.388 | 0.360 | DeepMSA2 |
| T1056 | 186 | 0.244 | 0.244 | qMSA |
| T1074 | 202 | 0.188 | 0.230 | q4JGI.a3m |
| T1082 | 97 | 0.488 | 0.488 | qMSA |
| T1090 | 193 | 0.402 | 0.463 | DeepJGI.hmsa3m |
| Avg | - | 0.388 | 0.401 | - |

Table **S4**: Improvement from bootstrapping on a randomly selected protein targets in the CASP13 dataset. Each sequence alignment (SA) set for a target contains 100 randomly generated sub-SAs. True lDDT scores of the original SA (before bootstrapping) and the true lDDT score of the top sub-SA (out of 100) as well as the best among the top-5 sub-SA are reported for comparison with percentage improvement in parenthesis.

| Target | SA Set | lDDT score of | | |
|---|---|---|---|---|
| | | Original SA | Top-ranked sub-SA | Best of top-5 sub-SAs |
| T1015s1 | uce0 | 0.573 | 0.587 (2.5%) | 0.589 (2.79%) |
| | uce3 | 0.607 | 0.564 (-7.14%) | 0.611 (0.64%) |
| | ure3 | 0.617 | 0.57 (-7.7%) | 0.62 (0.37%) |
| | ure5 | 0.622 | 0.617 (-0.95%) | 0.617 (-0.95%)) |
| T0980s1 | uce0 | 0.261 | 0.288 (10.24%) | 0.288 (10.24%) |
| | uce3 | 0.236 | 0.246 (3.98%) | 0.246 (3.98%) |
| | ure3 | 0.283 | 0.296 (4.78%) | 0.297 (5.06%) |
| | ure5 | 0.314 | 0.324 (3.25%) | 0.324 (3.25%) |
| T0968s2 | uce0 | 0.476 | 0.53 (11.25%) | 0.53 (11.25%) |
| | uce3 | 0.486 | 0.517 (6.34%) | 0.53 (9.08%) |
| | ure3 | 0.448 | 0.459 (2.28%) | 0.465 (3.73%) |
| | ure5 | 0.438 | 0.461 (5.11%) | 0.469 (6.98%) |
| T0968s1 | uce0 | 0.453 | 0.451 (-0.38%) | 0.472 (4.31%) |
| | uce3 | 0.339 | 0.339 (-0.09%) | 0.35 (3.18%) |
| | ure3 | 0.399 | 0.388 (-2.78%) | 0.404 (1.13%) |
| | ure5 | 0.392 | 0.381 (-2.91%) | 0.405 (3.14%) |
| T1017s2 | uce0 | 0.347 | 0.363 (4.73%) | 0.372 (7.18%) |
| | uce3 | 0.359 | 0.36 (0.33%) | 0.382 (6.35%) |
| | ure3 | 0.360 | 0.381 (5.69%) | 0.381 (5.69%) |
| | ure5 | 0.378 | 0.392 (3.62%) | 0.392 (3.62%) |
| T1001 | uce0 | 0.108 | 0.106 (-2.68%) | 0.112 (3.14%) |
| | uce3 | 0.159 | 0.159 (0%) | 0.159 (0%) |
| | ure3 | 0.334 | 0.35 (4.78%) | 0.356 (6.49%) |
| | ure5 | 0.240 | 0.253 (5.47%) | 0.253 (5.47%) |
| T0986s2 | uce0 | 0.403 | 0.403 (-0.12%) | 0.409 (1.44%) |
| | uce3 | 0.389 | 0.397 (2.03%) | 0.401 (3.29%) |
| | ure3 | 0.395 | 0.386 (-2.31%) | 0.404 (2.23%) |
| | ure5 | 0.395 | 0.384 (-2.79%) | 0.403 (2.15%) |
| T0957s1 | uce0 | 0.275 | 0.29 (5.34%) | 0.305 (10.76%) |
| | uce3 | 0.285 | 0.277 (-2.91%) | 0.287 (0.46%) |
| | ure3 | 0.326 | 0.329 (0.8%) | 0.329 (0.8%) |
| | ure5 | 0.260 | 0.257 (-0.92%) | 0.26 (0%) |
| T0957s2 | uce0 | 0.526 | 0.566 (7.76%) | 0.566 (7.76%) |
| | uce3 | 0.514 | 0.546 (6.31%) | 0.546 (6.31%) |
| | ure3 | 0.542 | 0.54 (-0.37%) | 0.549 (1.29%) |
| | ure5 | 0.529 | 0.53 (0.17%) | 0.542 (2.38%) |
| T1010 | uce0 | 0.116 | 0.116 (0.35%) | 0.116 (0.69%) |

| | | | | |
|---|---|---|---|---|
| | uce3 | 0.089 | 0.091 (2.25%) | 0.091 (2.25%) |
| | ure3 | 0.162 | 0.182 (12.68%) | 0.182 (12.68%) |
| | ure5 | 0.183 | 0.18 (-1.75%) | 0.189 (3.27%) |
| T0989 | uce0 | 0.280 | 0.28 (0.11%) | 0.289 (3.03%) |
| | uce3 | 0.236 | 0.239 (1.36%) | 0.249 (5.86%) |
| | ure3 | 0.226 | 0.225 (-0.44%) | 0.225 (-0.27%) |
| | ure5 | 0.213 | 0.213 (-0.23%) | 0.213 (-0.23%) |
| T0953s2 | uce0 | 0.193 | 0.192 (-0.62%) | 0.195 (1.03%) |
| | uce3 | 0.203 | 0.198 (-2.22%) | 0.204 (0.89%) |
| | ure3 | 0.253 | 0.253 (0%) | 0.253 (0%) |
| | ure5 | 0.250 | 0.251 (0.28%) | 0.251 (0.28%) |
| T0975 | uce0 | 0.204 | 0.203 (-0.49%) | 0.209 (2.45%) |
| | uce3 | 0.188 | 0.177 (-5.91%) | 0.188 (-0.11%) |
| | ure3 | 0.192 | 0.187 (-2.6%) | 0.187 (-2.6%) |
| | ure5 | 0.367 | 0.36 (-1.77%) | 0.36 (-1.77%) |
| T0950 | uce0 | 0.334 | 0.351 (5.15%) | 0.351 (5.15%) |
| | uce3 | 0.330 | 0.354 (7.06%) | 0.358 (8.3%) |
| | ure3 | 0.403 | 0.405 (0.42%) | 0.407 (0.94%) |
| | ure5 | 0.395 | 0.418 (5.8%) | 0.418 (5.8%) |
| T0990 | uce0 | 0.268 | 0.26 (-3.17%) | 0.268 (0%) |
| | uce3 | 0.263 | 0.263 (0%) | 0.273 (3.84%) |
| | ure3 | 0.359 | 0.359 (0%) | 0.359 (0%) |
| | ure5 | 0.359 | 0.27 (-24.71%) | 0.376 (4.79%) |
